# Mechanistic Origins of Dynamic Instability in Filaments from the Phage Tubulin, PhuZ

**DOI:** 10.1101/311498

**Authors:** Elena A. Zehr, Alexis Rohu, Yanxin Liu, Kliment A. Verba, Joe Pogliano, Nikolaus Grigorieff, David A. Agard

**Affiliations:** Department of Biochemistry and Biophysics and Howard Hughes Medical Institute, University of California, San Francisco, San Francisco, CA 94158, USA; Janelia Research Campus, Howard Hughes Medical Institute, 19700 Helix Drive, Ashburn, VA 20147, USA; Division of Biological Sciences, University of California, San Diego, CA 92093

## Abstract

A bacteriophage-encoded tubulin homologue, PhuZ, harnesses dynamic instability to position genomes of ՓKZ-like bacteriophage at the midline of their *Pseudomonas* hosts, facilitating phage infectivity. While much has been learned about molecular origins of microtubule dynamics, how GTP binding and hydrolysis control dynamics in the divergent 3-stranded PhuZ filaments is not understood. Here we present cryo-EM reconstructions of the PhuZ filament in a pre-hydrolysis (3.5Å) and three post-hydrolysis states (4.2 Å, 7.3 Å and 8.1 Å resolutions), likely representing distinct depolymerization stages. Core polymerization-induced structural changes reveal similarities to αβ-tubulin, suggesting broad conservation within the tubulin family. By contrast, GTP hydrolysis is sensed quite differently and is communicated by the divergent PhuZ C-terminus to the lateral interface, leading to PhuZ polymer destabilization. This provides a contrasting molecular description of how nucleotide state can be harnessed by the tubulin fold to regulate filament assembly, metastability and disassembly.

## INTRODUCTION

The tubulin cytoskeleton is an elaborate network of filaments that scaffolds and facilitates the transport of cellular contents. Microtubules harness the energy of nucleotide binding and hydrolysis to control their polymerization state and are notable in that they can stochastically switch between growth and disassembly phases, a process known as dynamic instability (Mitchison and Kirschner, 1984). In conjunction with a broad array of cellular proteins, this permits the spatial and temporal dynamics of the microtubule cytoskeleton to be exquisitely tuned for biological function. Underlying the dynamic instability phenomenon is that GTP hydrolysis is slow compared to filament growth allowing an intrinsically labile GDP core to be stabilized by GTP-bound subunits at its plus end (GTP cap) (Mitchison and Kirschner, 1984). The microtubule rapidly depolymerizes when hydrolysis overtakes cap growth, resulting in a process known as catastrophe.

Although the phenomenon has been studied for decades, the structural basis of dynamic instability is only recently being revealed by high-resolution cryo-EM reconstructions of mammalian microtubules in GTP-like and GDP states(Alushin et al., 2014; Zhang et al., 2015). These studies suggested that hydrolysis leads to lattice compaction and that the energy of GTP hydrolysis is stored in the energetically unfavorable configuration of the post-hydrolysis nucleotide binding site, the activation domain and H7 in α-tubulin. How general these findings are is as yet unclear; for example, the yeast microtubule does not appear to compact post-hydrolysis(Howes et al., 2017), suggesting that hydrolysis-induced strain can be built up without obvious structural alterations.

While dynamic instability was once thought to be a unique feature of microtubules, recently a bacteriophage-encoded tubulin homologue, called PhuZ (Kraemer et al., 2012) was found to also exhibit remarkably microtubule-like dynamic properties both *in vitro* and *in vivo* (Erb et al., 2014). The PhuZ family of tubulins is encoded by large ՓKZ-like *Pseudomonas* bacteriophages (Kraemer et al., 2012; Krylov et al., 2007). PhuZ filaments are structurally and biophysically polar and in the infected host cell, they are spatially organized such that their minus ends are anchored at the poles, and their plus ends radiate towards the cell center resembling a primitive spindle apparatus(Erb et al., 2014; Zehr et al., 2014). These dynamic spindles organize the bacteriophage 201Փ2-1 nucleoid, contained within a proteinaceous shell (Chaikeeratisak et al., 2017) at the cell midpoint, which by an unknown mechanism facilitates phage production(Kraemer et al., 2012). Recent observations suggest that in addition to nucleoid centering, empty capsids track along the filaments to facilitate DNA packaging at the shell surface (Chaikeeratisak et al., 2017).

Although primary sequence identity between PhuZ and αβ-tubulin is remarkably low (~15%), they share the same basic tubulin/FtsZ fold (Nogales et al., 1998). This consists of the two major functional domains: the GTP-binding domain (β-strands S1-S6 and α-helices H1-H5) and the GTPase activation domain (S7-S10 strands and H8-H10 helices) (Kraemer et al., 2012). The two domains are separated by the long helix H7, however, the otherwise conserved α-helix H6 is missing in PhuZ (Kraemer et al., 2012). The GTP-binding domain has a number of conserved loops that engage nucleotide (T1 through T6), and the activation domain has the T7 loop and helix H8 that stimulate hydrolysis of GTP via a set of conserved acidic residues (Kraemer et al., 2012; Nogales et al., 1998; Zhang et al., 2015).

As a consequence of significant sequence divergence beyond the core, PhuZ proteins form distinct higher-order assemblies with unique physical properties and ultrastructures. While PhuZ subunits within a protofilament are arranged in a canonical head-to-tail manner, the protofilament itself is structurally elastic unlike those of other tubulin homologues. The elasticity derives from a long and flexible C-terminus that provides extensive non-canonical longitudinal contacts, thereby stabilizing protofilament formation independent of precise head-to-tail positioning of adjacent subunits (Aylett et al., 2013; Kraemer et al., 2012). The C-termini also seem to function as flexible tethers allowing the protofilament longitudinal interfaces to compact and relax. Moderate resolution (7.1Å) cryo-EM studies of GMPCPP-stabilized filaments indicated that rather than being on the outside as in microtubules, the PhuZ C-termini face inwards to guide the assembly of the atypical three-stranded polymers through distinct lateral contacts to the three adjacent subunits (Zehr et al., 2014).

A comparison of X-ray crystallographic structures of PhuZ from the bacteriophage 201Փ2-1 in its soluble states (Aylett et al., 2013; Kraemer et al., 2012) with the polymer conformation derived from the cryo-EM map suggested a plausible assembly pathway (Zehr et al., 2014). Specifically, we proposed that GTP binding facilitates a compaction of the longitudinal interface from the unusually extended conformation seen in the crystal structure (Kraemer et al., 2012) into a more canonical tubulin/FtsZ-like longitudinal packing(Aylett et al., 2013). Upon assembly into filaments, the relatively straight starting protofilament twists, presumably making the filament lattice more GTPase-active and properly positioning the filament lateral surfaces to form assembly contacts. Concomitantly, the C-termini adopt a bent conformation, trading some of the longitudinal stabilization in order to form the filamentous lateral interactions. These observations suggested that the PhuZ filament might store the energy of GTP binding in the displacement of the C-terminal interactions and the supertwist of the filament (Zehr et al., 2014). At the medium resolution of the previous cryo-EM reconstruction we were able to infer only large-scale structural changes.

To address the central question of how GTP regulates PhuZ filament dynamic instability, in this work we focused on the GTP-to-GDP transition by using high-resolution cryo-EM to determine the structures of the PhuZ filament from bacteriophage ՓPA3 in both pre- and posthydrolysis states. We solved PhuZ in a GTP-like state (bound to GMPCPP) to ~3.5 Å and three different GDP states (4.2 Å, 7.3Å, 8.1 Å). The derived atomic model of the GMPCPP filaments more precisely elucidated polymerization-dependent changes in the PhuZ subunit, providing a more refined view of the polymerization mechanism. Additionally, we propose that the reconstructions of PhuZ-GDP filament represent different temporal states of its metastable lattice in the process of disassembly. Using a newly-implemented algorithm that determines local helical geometry parameters for each filament segment, we observe a break down of the helical order within the GDP-liganded lattice. These global changes are accompanied by local rearrangements in the T3 loop and C-terminus. The cryo-EM results in combination with negative stain EM analyses of the morphological changes in the disassembling PhuZ lattice, lead to a model wherein hydrolysis-triggered changes in the lateral filament interfaces that disorder the centrally-positioned C-terminal helix and cause the three-stranded helical lattice to progressively unwind into a highly-unstable state with a twisted ribbon architecture.

## RESULTS

### 3D Reconstruction of the PhuZ filament

We used cryo-EM to obtain a detailed view of the PhuZ filament lattice in a pre-hydrolysis state. Images were collected from PhuZ filaments polymerized with the non-hydrolyzable GTP analog, GMPCPP using a Polara microscope and K2 camera (Fig. 1A), and individual movie frames aligned to compensate for the beam-induced motion (Grant and Grigorieff, 2015; Li et al., 2013). Fourier transforms of aligned images illustrate the quality of the collected data with visible Thon rings extending to ~3.6 Å (Fig. 1A,inset).

**FIGURE 1.**
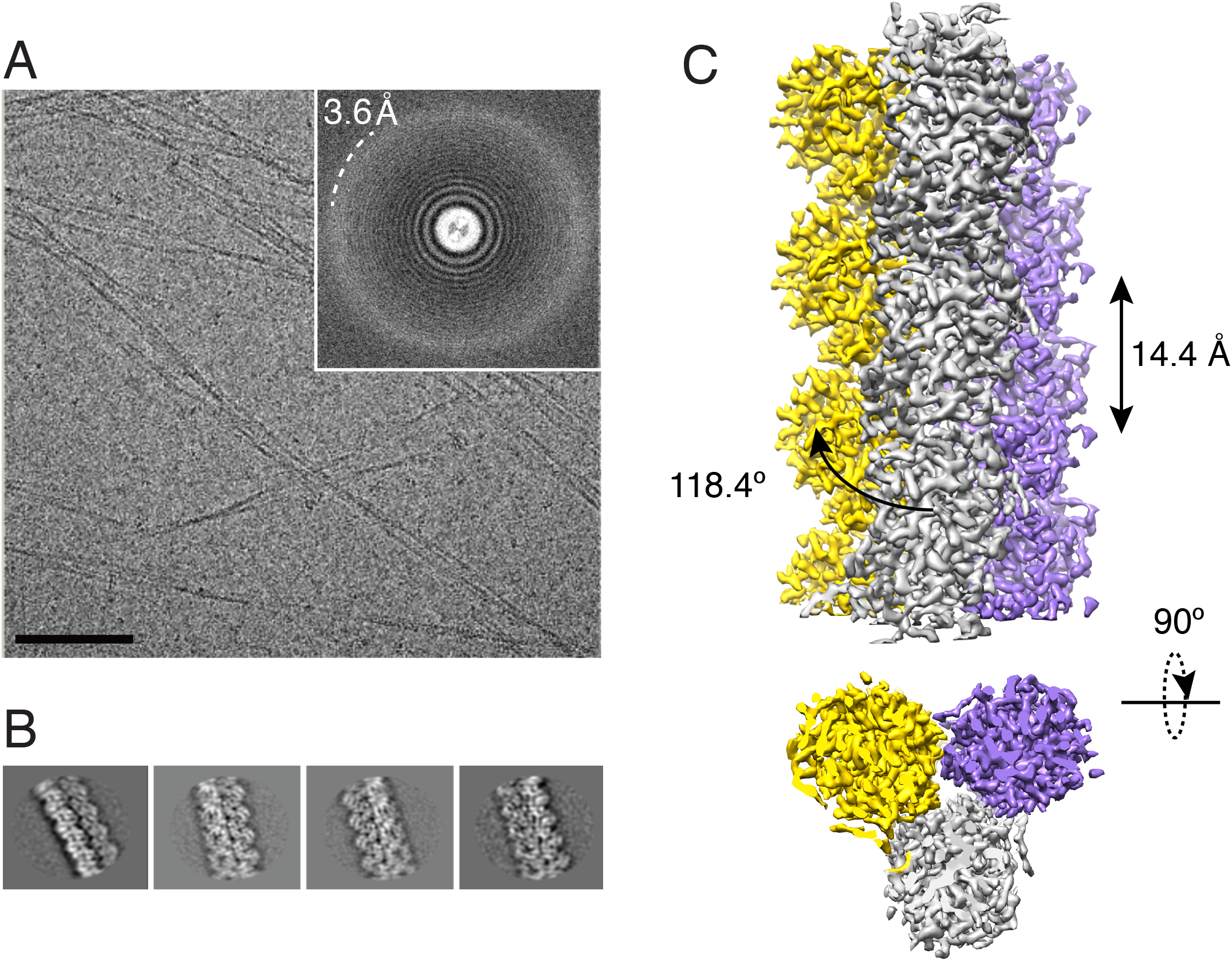
Data collection and 2D analysis on frozen-hydrated PhuZ filaments. **(A)** A micrograph of frozen-hydrated PhuZ filaments polymerized in GMPCPP (scale bar = 100 nm) and its power spectrum with tone rings extending to 3.2 Å (inset). **(B)** Reference-free 2D class averages of PhuZ-GMPCPP filament segments show secondary structure elements. **(C)** 3.5Å Cryo-EM reconstruction of PhuZ filament in GTP-like state. The filament is a left-handed one-start helix, with an overall right-handed supertwist.

We used extensive image sorting in combination with single-particle helical analysis methods to obtain an initial 3D reconstruction of PhuZ filament (Materials and Methods). First, to get a homogeneous subset of filament segments, the data were extensively classified in 2D with RELION (Scheres, 2012). Class averages revealed clearly discernable secondary structure elements and suggested high structural order of the PhuZ-GMPCPP filament lattice (Fig. 1B). An initial reconstruction at 4.5 Å (FSC=0.143, with fully independent half datasets) was obtained using iterative helical real space reconstruction (IHRSR) (Table S1, Fig. S1A) (Egelman, 2000, 2007; Frank et al., 1996; Scheres and Chen, 2012). IHRSR-estimated helical symmetry parameters were 118.4° rotation and 14.4 Å axial translation per subunit, a helical geometry slightly different from its previously studied homologue (Zehr et al., 2014). Filament segments were further classified in 3D against multiple references and their parameters were refined with FREALIGN (Grigorieff, 2007) (Table S1, Fig. S1A). Finally, the map was refined by re-estimating movie whole-frame shifts using Unblur (Grant and Grigorieff, 2015) and CTF parameters using CTFFIND4 (Rohou and Grigorieff, 2015). Alignment parameters for filament segments were refined using Frealix (Rohou and Grigorieff, 2014) operated in a single-particle mode, and modified to compensate for the beam-induced motion local to each filament segment (details in Methods).

The nominal resolution of the final reconstruction (Fig. 1C, S1a) was 3.5Å (FSC=0.143 cutoff, Table S1, Fig. S1A). Details consistent with this resolution estimate are evident and distributed more uniformly throughout the subunit in the Frealix reconstruction than with either of the other two methods. This improved uniformity of the resolution (Fig. 2) is also reflected in the much sharper falloff in the FSC curve (Fig. S1A). Individual β-strands are clearly separated, the majority of side-chains are visible (Fig. 2B,C), and the nucleotide-binding site, including densities for GMPCPP and the Mg^2^+ ion are well resolved (Fig. 2A). To fully interpret high-resolution features of the map, we built an atomic model of PhuZ-CMPCPP (details in Methods).

**FIGURE 2.**
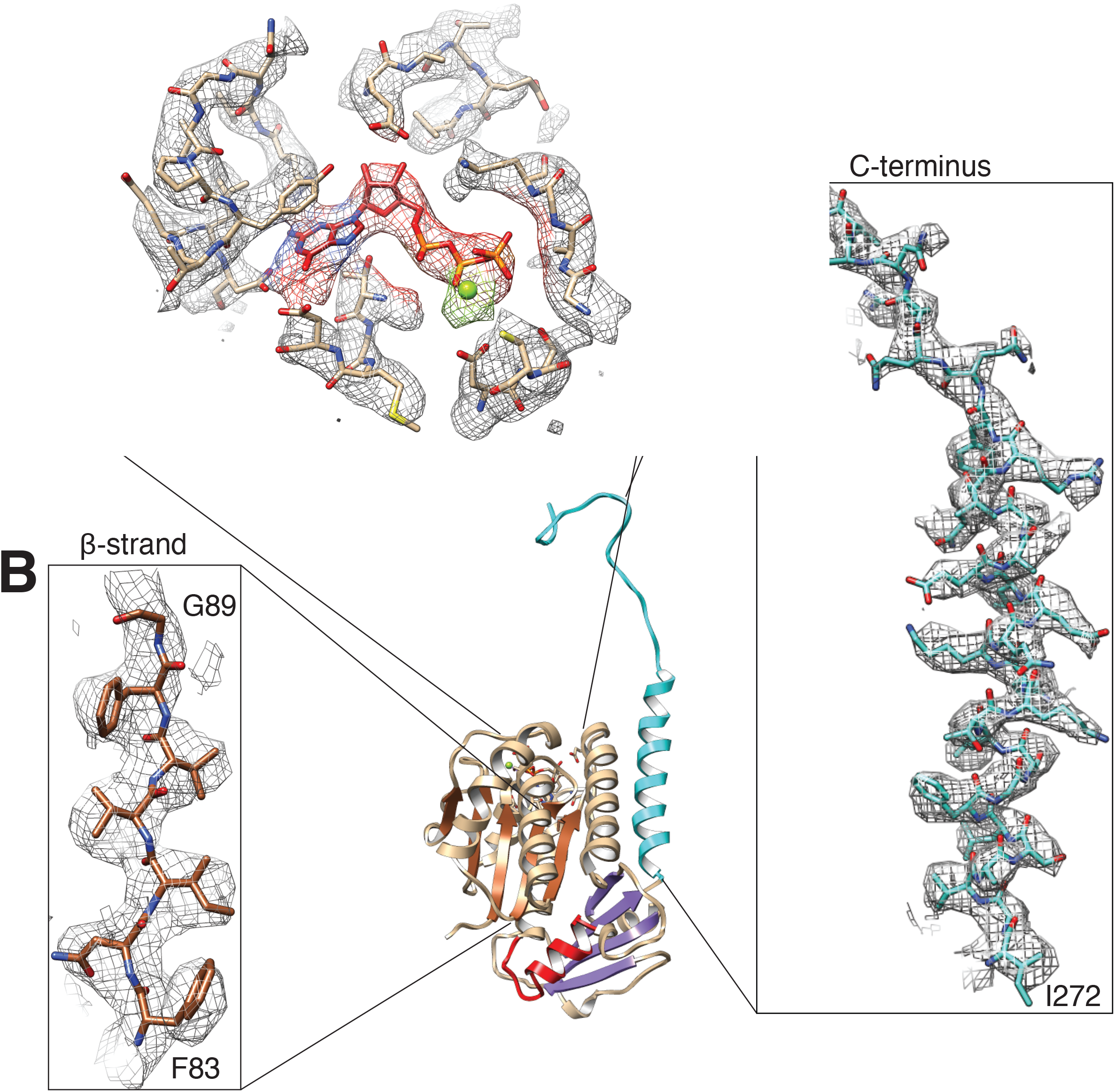
Atomic-level 3D view of PhuZ-GMPCPP filament. 3D density map of PhuZ-GMPCPP filament filtered to 3.5 Å resolution shown as mesh isosurfaces **(A-C)**. A ribbon diagram of the atomic model of PhuZ bound to GTP derived from the map **(D)**. **(A)** The GTP-binding site with the loops T2-T6 and helices H1 and H7 that bind GTP, colored in red. Mg^2+^ and its corresponding EM density are colored in green. **(B)** The beta strand S4 (residues 83 - 89) found within the GTP-binding domain of PhuZ shows well-resolved EM densities for all of the side-chains. **(C)** A portion of the C-terminus (displayed residues 272 - 302 of the total 272 - 315) with most of the side-chains seen in the 3D reconstruction.

### Polymerization is accompanied by alterations of the H7 helix, the activation domain and the longitudinal interface

In addition to the previously-reported large-scale rearrangements of the C-terminus upon polymerization (Zehr et al., 2014), we can now resolve a restructuring of the PhuZ longitudinal interface and alterations within its tubulin fold. For comparison, the two longitudinal dimers representing a solution state of PhuZ (PDB ID: 3r4v) (Kraemer et al., 2012) and its polymer form (this work) were aligned by the β-sheet of the GTP-binding domain, excluding the short and most structurally-variable strand S3 (Fig. 3C). A morph between the dimers reveals a 1.2 Å longitudinal shift of the H7 towards the plus-end interface, a bending of the top of this helix, and a repositioning of the T6 loop by 2.2Å such that it establishes contacts with the H10-S9 loop of the subunit up above (Fig. 3A; Movie 1). Additionally, the motion in the top of H7 maintains the pi-stacking interactions between the nucleotide base and Y161 of H7, and hydrogen bonding of the base with D165, found one α-helical turn below (Fig. 3A; Movie 1). Similar interactions have been described for other tubulin homologues, such as α/β-tubulin and FtsZ (Lowe and Amos, 1998; Lowe et al., 2001).

**FIGURE 3.**
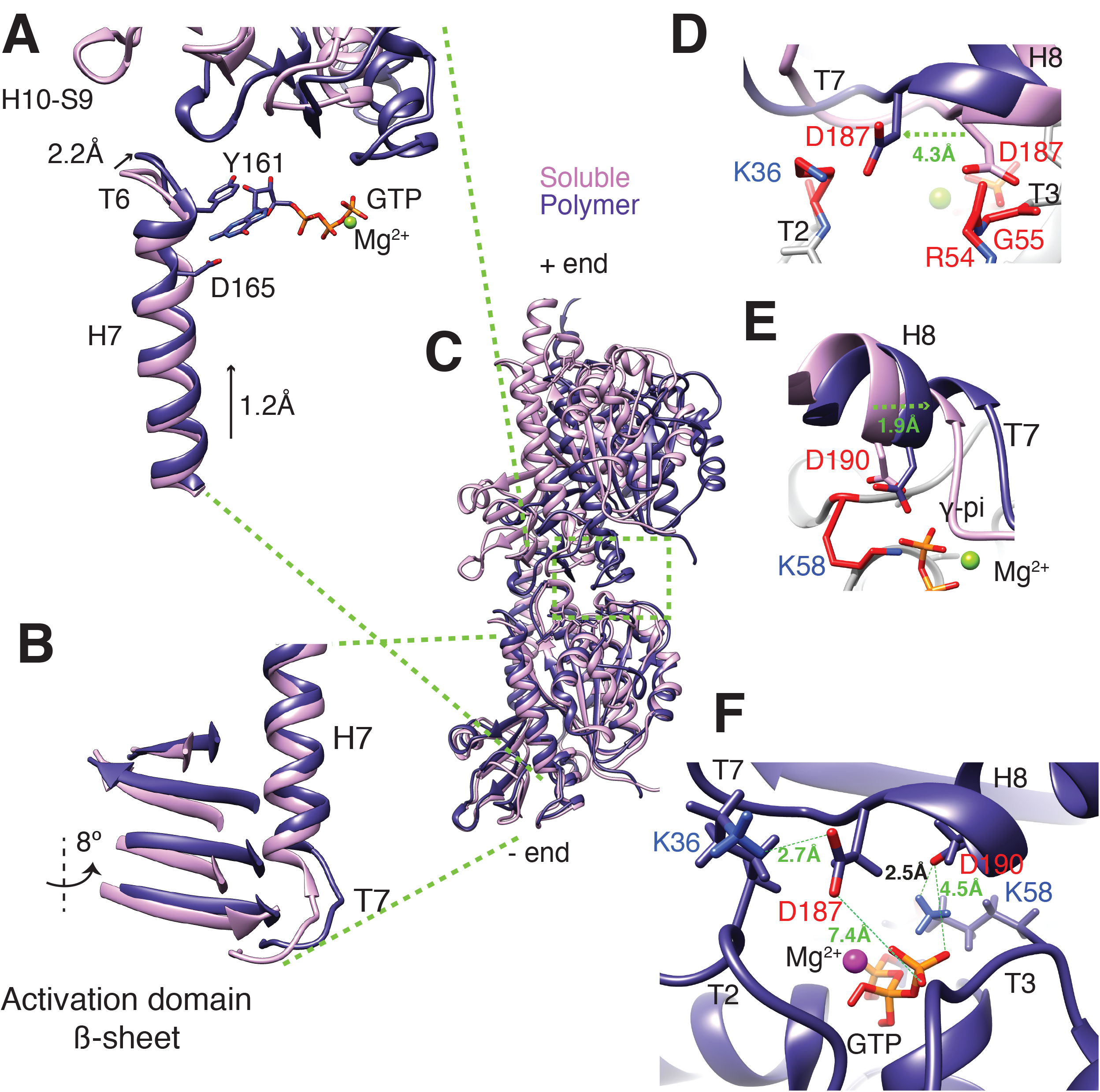
Structural changes that accompany PhuZ polymerization. **(A)** Ribbon diagram depicting the H7 and adjacent loop T6 of one subunit and a part of the activation domain of the longitudinally adjacent subunit including the H10-S9 loop in soluble (*violet*) and polymer states (*purple*). The T6 and H7 undergo subtle conformational changes, in response to compaction and twisting of the longitudinal interface. Residues Y161 and D165 that bind the base of GTP are shown as sticks. **(B)** A ribbon diagram showing the restructuring of the activation domain with only of its beta sheet displayed and T7 loop in response to the upward movement of the H7. **(D-F)** The GTPase site of PhuZ filament shows catalytic D187 (T7) and D190 (H8) making ionic bonds with the lysines 36 (T2) and 58 (T3), respectively, as deduced from the cryo-EM reconstruction and molecular dynamics calculations.

Helix 7 provides a mechanism to couple these changes at the longitudinal interface and nucleotide-binding site to alterations in the activation domain and the minus-end interface, which in turn interacts with the adjacent subunit’s plus-end interface (Fig. 3B; Movie 1). The side-chains for K36 (T2) at the minus end and the catalytic D187 of the subunit at the plus end, both well resolved in the map, are positioned to support a salt bridge (Fig. 3D,F). EM density for the other catalytic aspartate side chain, D190, is missing, but based on the fitting results combined with the MDFF molecular dynamics simulations, D190 could well be positioned to make ionic interactions with K58 in T3 (Fig. 3E,F).

Finally, the changes at the longitudinal interface are accompanied by an ~8° rotation of the activation domain in the direction of the GTP-binding domain. Helix 7 and T7 make a hydrophobic interface and form a number of hydrogen bonds with the activation domain. The net effect of all these motions is that the plus end, minus end and the activation domain all reorganize as a single unit in response to polymerization. This provides a tight coupling between conformational changes within the tubulin monomer, formation of an optimal longitudinal interface, and the activation of the catalytic machinery for GTP hydrolysis (Fig 3B; Movie 1).

### The lateral filament interface is formed by conserved residues

Previously, based on an pproximate main chain juxtaposition, we proposed that the lateral interface was stabilized by ionic interactions between conserved acidic residues in the C-tail (D303 and D305) and positively charged residues (R238 and R217, respectively) across the lateral interface (Zehr et al., 2014). This was tested *in vitro* and *in vivo* by making R217D and D305R mutations in PhuZ. While the single charge-swap mutations abrogated three-stranded filament formation, the double mutation partially restored polymerization, supporting the proposed interaction. The current reconstruction, has well-resolved densities corresponding to these side-chains allowing them to be precisely positioned, with the exception of D305. R217 is sandwiched between D303 and D305 and is positioned to form electrostatic interface with either of these aspartates, while R238 forms interface with D305 (Fig. S2A). This may explain why the double mutant R217D/D305R was unable to fully restore activity to wild-type levels (Zehr et al., 2014).

The better resolved structure also revealed additional lateral interactions. The two hydrophobic residues I304 and V240 reinforce interactions between the tail and H9/H10 (Fig. S2A). The helices 11 of the three strands make lateral contacts with each other in a staggered manner. H11 of one stand simultaneously contacts helices 11 of the two other strands via its C-terminal conserved E287 and R294, which explains the cooperative assembly behavior of PhuZ filaments (Fig. S2B).

### GTP hydrolysis and phosphate release lead to global rearrangements and filament disassembly

To understand molecular mechanisms underlying PhuZ dynamic instability and the role of GTP hydrolysis, we wanted to obtain a high-resolution filament structure for the PhuZ-GDP state. Because PhuZ, unlike many other tubulins, does not polymerize in high concentrations of GDP (Fig. S3D) (Kraemer et al., 2012), PhuZ filaments were assembled in low concentrations of GTP and then allowed to age in solution before visualization. Light scattering shows rapid polymer growth during the first 30 seconds following addition of 400 μM GTP to 40 μM PhuZ, followed by a slower rise to plateau at 150 to 300 seconds (Fig. 4A, inset). Whether this slower rise phase is due to increased bundling, changes in the filament morphology or decreased GTP levels is unclear. The scattering intensity started to decay at 400 seconds, suggesting conversion from net filament assembly to net disassembly.

**FIGURE 4.**
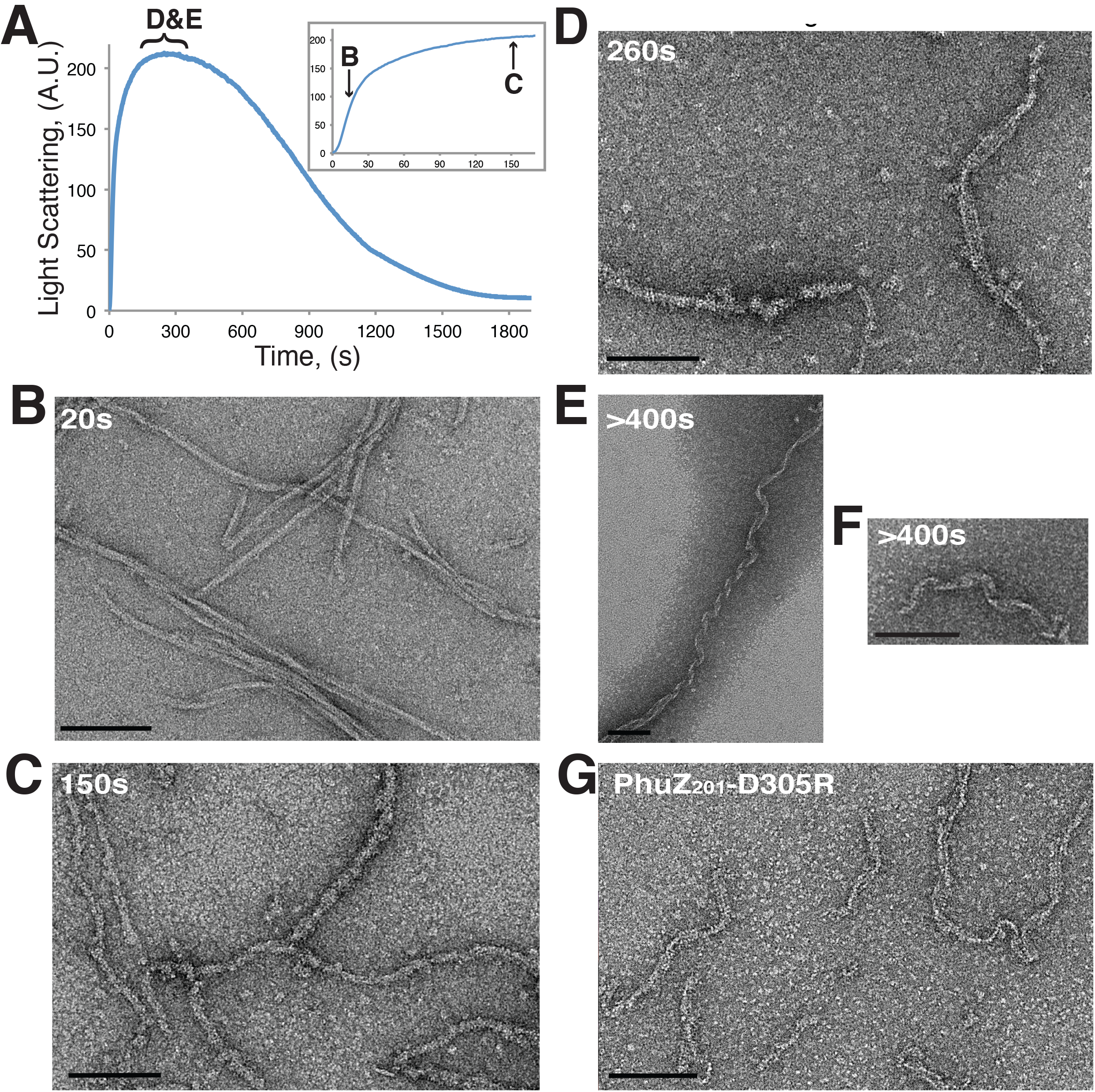
Negative-stain EM analysis of disassembling PhuZ filaments. **(A)** Right-angle light-scattering traces of 40 μM PhuZ polymerized in 400 μM GTP. PhuZ filament population was analyzed by negative-stain EM **(B-F)** at different time points after GTP addition. The time points are indicated on the curve in bold letters **(A)**, which correspond to the panel labeling in this figure **(B-F)**. **(A)** Negatively-strained PhuZ filaments are long, well-ordered and straight at 20 s post-GTP addition. **(B)** PhuZ filaments have rough appearance and high curvature when visualized at 150 s post-GTP addition. **(C)** The PhuZ lattice unwinds into flat sheets at 260 s after GTP addition and assumes twisted-ribbon architecture (**E** and **F**). PhuZ filaments with the twisted-ribbon morphology resemble a mutant in the lateral interface, D305R. This mutation was made in the PhuZ homologue from the bacteriophage 201Փ2-1 (PhuZ-D305R) and described in Zehr et al., 2014. (G) Scale bar = 100 nm.

Morphological changes of the PhuZ filament were assessed at different times after GTP addition by negative stain EM. During the rapid growth phase, filaments appeared to be long, well-ordered and fairly straight (Fig. 4B; Fig. S3A). This morphology was similar to that of GMPCPP filaments, suggesting that widespread hydrolysis had not yet occurred in the lattice (Fig. S3B). In contrast, the filaments exhibited a much rougher appearance with a non-uniform diameter and less distinct subunit organization at the onset of the equilibrium phase (Fig. 4C; Fig. S3C). To test whether this lattice roughening was triggered by GTP hydrolysis or Pi release, we imaged filaments polymerized in the presence of GTP and 1mM beryllium fluoride (BeF_3_). BeF_3_ is a phosphate analogue that mimics and stabilizes the post cleavage (ADP-Pi) transition intermediate formed after γ-phosphate cleavage, but before phosphate dissociation. Since these filaments were stable, quite uniform, and devoid of the rough appearance (Fig. S3E), we inferred that filament roughening and its subsequent disassembly were triggered not by GTP hydrolysis, but by the subsequent release of the cleaved phosphate. Moreover, the large difference in scattering amplitude between 100 *μ*M GTP and 100*μ*M GTP+BeF_3_ (Fig. S3D) implies that GTP hydrolysis and phosphate release leads to unproductive filament formation/disassembly even in the earliest phases of PhuZ polymerization.

In the late stages of the equilibrium phase the rough filament appeared unwound along its length, forming flat sheets of parallel protofilaments (Fig. 4D; Fig. S3F). Some of these sheets assumed a writhed-ribbon morphology when not physically restrained by the rough filament lattice at their ends (Fig. 4E,F). This writhed-ribbon architecture resembles that of the PhuZ lateral interface mutant D305R (Fig. 4G; Zehr et al., 2014), suggesting this interface becomes particularly labile following GTP hydrolysis and phosphate release. Owning to the heterogeneity of the twist in the ribbon-like structures, we were unable to obtain meaningful reference-free 2D averages to get a more detailed view of this architecture.

Taken together, these negative stain results combined with the BeF3 light scattering data established that hydrolysis occurs early during the assembly phase, and that the PhuZ-GDP lattice, presumably with a GTP cap, is fairly stable and continues to support growth for some time.

### GTP hydrolysis leads to increased helical disorder

To gain a molecular-level appreciation of the early effects of GTP hydrolysis, we sought to determine a cryo-EM reconstruction of the well-ordered, yet metastable, PhuZ-GDP filaments present in the steady-state phase. At various time intervals following GTP addition, we vitrified samples as previously for GMPCPP filaments. Vitrification 150 seconds after GTP addition was successful, but at later time points very few filaments could be observed, suggesting that the later disassembly intermediates seen by negative stain (Fig. 4) had been stabilized by the carbon support.

The cryo-EM reconstruction of the 150-second time point sample was carried out as described earlier, except that the dataset appeared significantly more heterogeneous, making it even more important to pursue 3D classification with FREALIGN. This yielded three distinct classes with a little more than half of the images grouped into classes 1, 2A and 2B (Fig. 5A,B, Table S3). Class 1 contained about 33% of the filament segments and after refinement in Frealix resulted in a 4.2 Å resolution map (Fig. S4A, Table S3). By contrast, the other two classes (classes 2A and 2B) yielded maps at considerably lower resolutions (8.1 Å and 7.3 Å respectively, Fig. S4A, Table S3) and displayed hollow or partially hollow lumens (Fig. 5A). This was somewhat surprising given that in the GMPCPP map the long helix H11 and the C-terminus, which together fill the lumen, were the best-ordered regions of the map. Attempts to obtain higher resolution reconstructions by combining classes 2A and 2B and further refinement in Frealix or classification in FREALIGN did not significantly improve the map (not shown).

**FIGURE 5.**
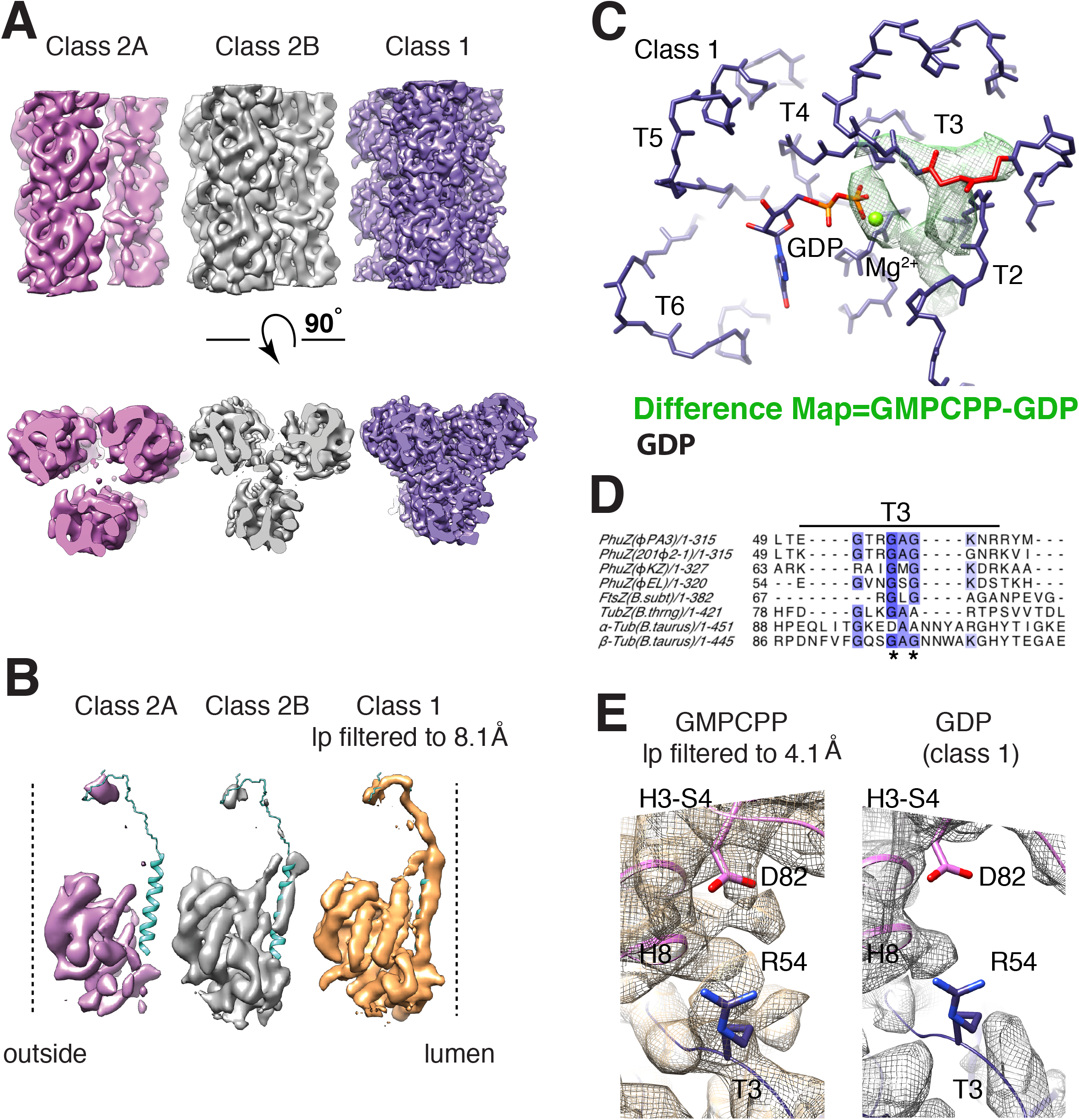
GDP-liganded lattice displays relaxation of the lateral and longitudinal interfaces. **(A)** Three 3D PhuZ-GDP classes isolated using FREALIGN. Class 1 is the highest resolution GDP class that shows only subtle differences from the GMPCPP structure, and the central lumen is completely filled. The classes 2A and 2B show hollow or partially hollow lumens. **(B)** The cryo-EM 3D classes of PhuZ-GDP, show relaxation of the C-terminus. A PhuZ subunit, extracted from each class, is shown as a solid isosurface, colored in *violet* for the class A and *gray* for the class B. The Class 1 of PhuZ-GDP (originally solved at 4.2 Å) was low-pass filtered to 8.1 Å for comparison and is shown as a *tan* solid isosurface. A ribbon diagram depicting the C-terminus is in *turquoise*. The outer surface of PhuZ filament is labeled as *outside* and the inner as *lumen*. **(C)** A detailed view of PhuZ GTP-binding site with the loops labeled as T2 -T6. Only the backbone atoms are displayed. R54 and G55 are shown in *red*, GTP is colored by heteroatom, the magnesium ion is in *green* and the rest of the atomic model is in *purple*. A calculated difference map between 3D reconstructions of PhuZ-GMPCPP and PhuZ-GDP (class 1) is shown as a *green* mesh isosurface. To generate the difference map both the PhuZ-GMPCPP and the class 1 maps were scaled against an atomic model, low-pass filtered to 4.2 Å with the Butterworth filter and subtracted from each other. For visualization purposes smaller disconnected parts of the difference map were hidden with the “hide dust” option using UCSF Chimera. The difference map shows density features around the γ-phosphate, R54 and G55. (D) Alignment of amino acid sequences of tubulin homologues using Mafft with default settings in Jalview (Waterhouse et al., 2009). The amino acid positions are indicated at the beginning of each line. Abbreviations for sequences encoding tubulin homologues are: PhuZ(201Φ2-1) from phage 201Φ2-1, PhuZ(ΦPA3) from phage ΦPA3, PhuZ(ΦKZ) from phage ΦKZ, PhuZ(ΦEL) from phage ΦEL, FtsZ from *Bacillus subtilis*, TubZ from *Bacillus thuringiensis* and α-and β-tubulin from *Bos Taurus*. Conserved sequences are highlighted in dark or light *blue* to show highly and less conserved residues, respectively. R54 and G55 are marked with (⋆). **(E)** D82 in the H3-S4 loop and R54 in the T3 loop make an ionic bond in PhuZ-GMPCPP reconstruction but not in PhuZ-GDP (class 1) map. Both maps were low-pass filtered to 4.2 Å for comparison.

Driven by the profound changes seen in negative stain, we were particularly concerned that helical disorder within the relaxing PhuZ lattice could be the dominant factor limiting resolution. The local helical parameters for each filament segment were estimated using Frealix for all datasets (GMPCPP, GDP Classes 1, 2A, 2B; Fig. S4B, Table 1,2). Taken together, these estimates indicated changes upon GTP hydrolysis both in the mean geometry of the helical lattice, which becomes more elongated, and in helical flexibility, which is significantly increased in the GDP dataset, especially for classes 2A,B. Attempts to improve map quality by omitting filament segments with helical parameters deviating beyond one standard deviation in the GMCPP and all 3 GDP classes (Table 2) had minimal effects on the maps, improving quality in a few areas (Fig. S5A,C), but decreasing it elsewhere. The unchanged resolution as reflected by the FSC (Fig. S5B,D) is likely due to the 40% decrease in segment number for both sets (Table 2).

**Table 1:** Helical Symmetry Parameters for PhuZ Filament Segments.

Twist per subunit
| Reconstruction | Mean, (deg) | SD, (deg) | 95% particles |
| --- | --- | --- | --- |
| GMPCPP | 118.39 | ±0.55 | 117.27-119.61 |
| GDP Class 1 | 118.66 | ±0.62 | 117.34-119.90 |
| GDP Class 2A | 118.72 | ±0.87 | 116.91– 120.41 |
| GDP Class 2B | 118.70 | ±0.85 | 116.96 –120.50 |

Rise per subunit
| Reconstruction | Mean, (deg) | SD, (deg) | 95% particles |
| --- | --- | --- | --- |
| GMPCPP | 14.60 | ±1.16 | 11.66-16.98 |
| GDP Class1 | 14.82 | ±1.18 | 12.74-17.57 |
| GDP Class A | 15.17 | ±1.24 | 13.44– 18.24 |
| GDP Class B | 15.15 | ± 1.23 | 13.32– 18.16 |

**Table 2:** Parameters for the PhuZ Maps Refined in FREALIX.

| Reconstruction | Total asym units | Selected asym units | Twist Range, (deg) | Rise range, (Å) | Resolution, (Å) |
| --- | --- | --- | --- | --- | --- |
| GMPCPP | 85,472 | 51,811 | 117.80-118.90 | 14.10-15.00 | 3.5 |
| GDP Class 1 | 68,147 | 41,432 | 118.20-119.10 | 14.30-15.00 | 4.2 |
| GDP Class 2A | 34,218 | 12,442 | 117.93-119.53 | 14.17-15.17 | 8.1 |
| GDP Class 2B | 31,772 | 12,403 | 117.92-119.52 | 14.09-15.09 | 7.3 |

The class 2 segments are the most structurally flexible, having widely varying twist and rise values (Fig. S2A and B, Table 1). Selecting segments from this class to limit the spread in helical parameters to 1.9° twist and 1.7Å rise, (Table 2) only marginally improved the overall visual quality of the reconstruction (Fig. S4E), and marginally decreased the estimated resolution from 7.3 Å to 8.1 Å (Table 2, Fig. S4F; note the two FSC curves falloff at exactly the same resolution). This is likely due to the quite severe 64% reduction in segment number. Significantly larger data sets and even more restrictive helical segment selection would probably be required to meaningfully extend the resolution of these classes.

### GTP hydrolysis weakens filament interfaces

The increased disorder in the GDP lattice is likely a direct consequence of GTP-hydrolysis-induced changes in the intra- and interprotofilament interfaces. The missing luminal density (Fig 5A) in the classes 2A and 2B corresponds to the PhuZ C-terminus and suggests its relaxation post hydrolysis (Fig. 5B). In class 2A the C-terminal density is missing except for the very end of the C-terminal tail, residues ~ 306 – 315 of the “knuckle” (Kraemer et al., 2012). The C-terminus in class 2B is more ordered: in addition to the ten most C-terminal residues, weak density corresponding to H11 is also present (Fig. 5B). As shown in the crystal structure, the negatively charged knuckle of one subunit interacts strongly with a basic patch, formed by helixes H3-H5, of its longitudinal neighbor (Kraemer et al., 2012). This interaction also remains intact during PhuZ polymerization (Zehr et al., 2014), even though the rest of the C-terminal structure rearranges. That only the core of the knuckle region remains intact in classes 2A and 2B is surprising. suggesting that not only are H11 and the tail disordered, but also the basic patch with which it interacts. Additionally, a number of loops facing the lumen are also not resolved. Among these are the S6-H7 and H9-S8 loops, which form inter-protofilament contacts with the C-terminus (Zehr et al., 2014). These cryo-EM results are consistent with the observations on the negatively stained disassembling filaments displaying relaxation between the strands.

The much higher resolution of GDP class 1 permits a more detailed analysis of local changes, likely corresponding to the earliest phase of post-hydrolysis disassembly. Density for the γ-phosphate and Mg^2^+ are greatly reduced as observed by strong negative densities in a GMPCPP-Class 1 difference map (Fig. 5C). Particularly notable is a structural reorganization of T3 loop, where density for residues G55 and R54 are also greatly reduced (Fig. 5E). G55 is part of the conserved tubulin GXG motif (Fig. 5D) that in the GMPCPP structure contacts the γ-phosphate, (Fig. 2A, Fig. 5C,E). Generally, loss of the γ-phosphate causes relaxation of the loop. Indeed, the T3 loop is frequently disordered in crystal structures of tubulin homologues bound to GDP (Aylett et al., 2013; Kraemer et al., 2012). This is important as the loop appears to form longitudinally-stabilizing interactions in numerous tubulin-like protofilaments as well as our PhuZ. In PhuZ:GMPCPP, T3 loop residues R54 and K58 form salt bridges with D82 (Fig. 5E) and D190 (Fig. 3C,D) of the next monomer, respectively. In the GDP Class 1 map, the R54-D82 density is completely missing (Fig. 5E), indicating a weakening of the post-hydrolysis longitudinal interface. To verify this observation, we made the D82A mutant, and it failed to polymerize in excess GMPCPP (data not shown).

## DISCUSSION

It has long been appreciated that NTPs facilitate assembly of actins and tubulins into polymers, and consequently, nucleotide hydrolysis favors disassembly. While the process is common, very few polymers exhibit dynamic instability – that is the rapid switch between polar growth and catastrophic depolymerization. For this to happen requires the ability to store a significant amount of strain energy within the filament lattice and to have a sufficiently stable cap can persist even after the bulk of the filament has hydrolyzed NTP, switching into a trapped metastable NDP state. The best studied of dynamically unstable polymers is the heterodimeric eukaryotic tubulin, which forms 13 protofilament microtubules. Advances in cryoEM have led to high-resolution structures of mammalian and yeast microtubules in GTP-like states (refs) revealing how the solution form converts from a curved geometry to being straight in the microtubule lattice, storing energy in the process (Luke PNAS). However, only subtle changes where observed in the lattice upon GTP hydrolysis in mammalian tubulin (ref) and no changes were seen with yeast microtubules (ref). The fundamental challenge is that lattice cooperativity seeks to minimize observable changes. Here we provide new insights into how the energy of GTP binding can be captured and released by the only other dynamically unstable tubulin, the highly divergent monomeric bacteriophage PhuZ.

To get insight into the sequence of conformational changes during PhuZ turnover, this work extends our previous understanding of PhuZ polymerization (Zehr et al., 2014), and examines the filament’s disassembly mechanism. Using cryo-EM and a new image-processing algorithm (Rohou and Grigorieff, 2014), we capture the near-atomic structure of the PhuZ filament in its GTP state and describe its polymerization cycle in finer details. We also explain structural changes in PhuZ filament as it progresses through the early stages of disassembly.

We propose that metastability of the PhuZ lattice is stored in the twist of the PhuZ protofilaments, the large-scale rearrangements of the PhuZ C-terminus (Zehr et al., 2014), the energetically unfavorable displacements of the H7 and the activation domain. This energy is released as the γ-phosphate-sensing T3 loop relaxes, and the key longitudinal interactions disappear (Fig. 6). Weakening of the longitudinal interface allows each PhuZ subunit to rotate and translate more freely about the protofilaments’ central axes, which is reflected in the wide spread of the twist and rise per subunit values for the GDP segments. Subsequently, the PhuZ filament loses its lateral contacts mediated by the C-terminus. The progressive accumulation of the helical lattice disorder leads to the global restructuring of the filament’s architecture from three-stranded to the writhed filamens seen in negative stain.

**FIGURE 6.**
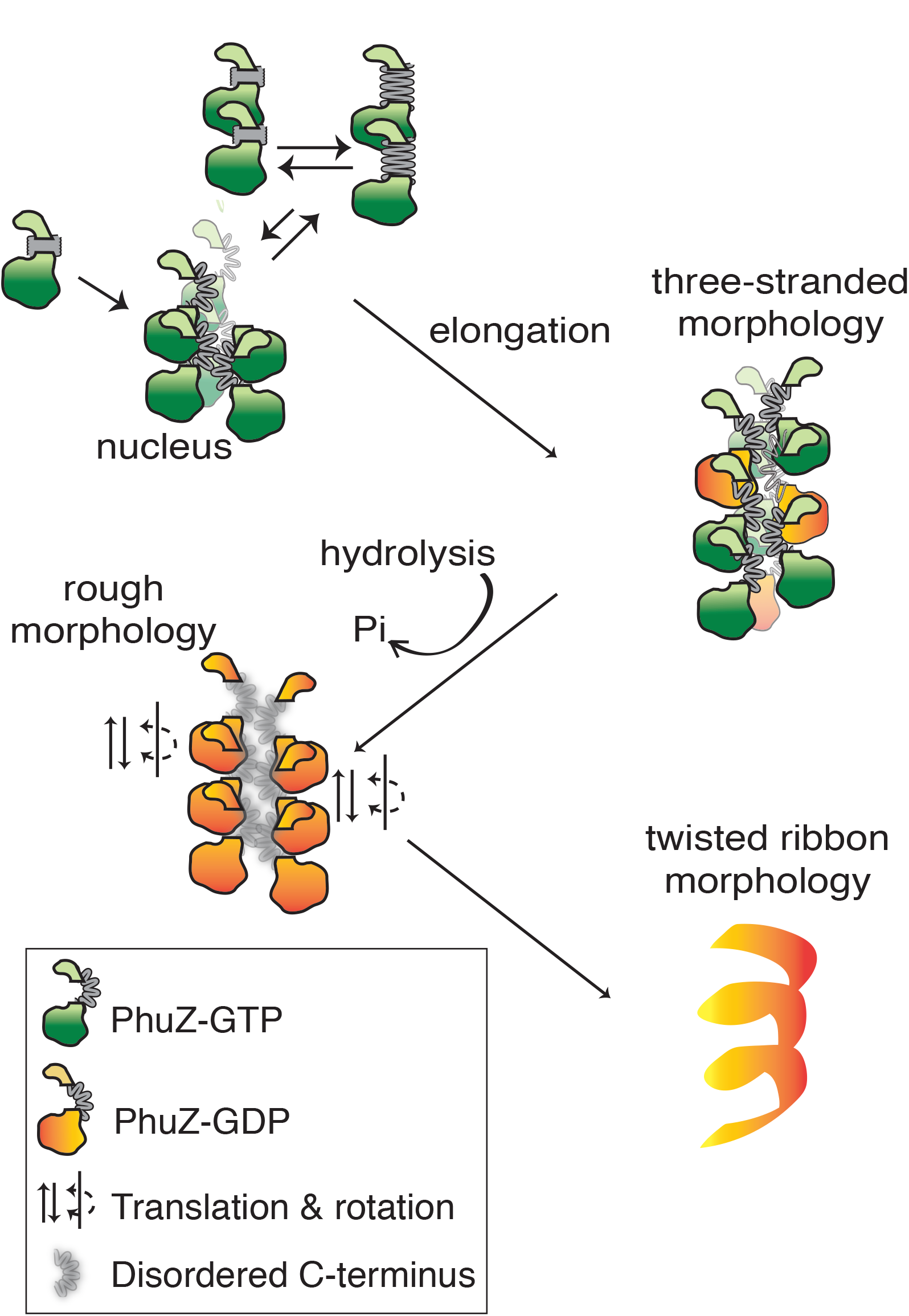
Refined model for PhuZ assembly/disassembly cycle. PhuZ monomers assemble dimers with the tense longitudinal interface as in the crystal structure (PDB ID: 3ZBQ) (Aylett et al., 2013) or relaxed interface as in the crystal structure (PDB ID: 3r4v) (Kraemer et al., 2012). Three dimers form a hexameric nucleus with subunits displaying the filament contacts as described in (Zehr et al., 2014) and this work. Polymerization is accompanied by rearrangements of the C-terminus (Zehr et al., 2014), H7 and the activation domain. The nucleus grows by the addition of dimers and monomers. GTP is hydrolyzed soon after filament assembly leading to relaxation of longitudinal (the T3 loop) and lateral (the C-terminus) contacts. The PhuZ lattice unwinds into sheets that supercoil into fragile polymers with twisted ribbon architectures.

Due to increased structural disorder and fragility of the PhuZ filament lattice post-GTP hydrolysis, we were unable to resolve the GDP-bound lattice to a near atomic view. In the future, it would be necessary to stabilize the disassembling lattice by, for example, capping the filament’s ends with GTP-bound subunits. Although at low-resolutions the PhuZ lattice looks highly ordered when bound to BeF_3_, it will be essential to obtain a near-atomic structure of the filament’s restructuring GTP-binding site in response to hydrolysis.

## MATERIALS AND METHODS

### Protein Expression and Purification

The gene encoding PhuZ from ՓPA3 was cloned into pET2w8a (+) with a 6xHis-tag on the N-terminus and expressed in BL21(DE3) cells under an IPTG-inducible T7 promoter. 1 mM IPTG was added once cells reached an OD_600_ of 0.7 at 37°C and protein was allowed to express for 8 hr at 16°C before the cells were pelleted. Cells were lysed in a buffer containing 250 mM KCl, 50 mM HEPES pH 7.2, 1 mM MgCl_2_, 10% glycerol and 1mM DTT. EDTA-free protease inhibitor tablets were included during lysis. 250 mM imidazole was added to elute the protein from the Ni-NTA resin. The 6xHis-tag was cleaved overnight at 4°C by thrombin protease, followed by gel filtration chromatography (Superdex 200) in BRB80 pH 7.2, 1mM DTT.

### Specimen preparation

#### PhuZ-GTP cryo-EM sample preparation

PhuZ was thawed and spun at 80,000X RPM for 20 min in a TLA-100 rotor (Beckman) at 4° C to remove protein aggregates. 50 μM PhuZ was polymerized in BRB80 pH7.2 and 1 mM GMPCPP for 1 minute at room temperature. 2 μl samples were applied to glow-discharged C-FLAT holey carbon grids with 1.2μm hole size, 400 mesh. The grid was blotted for 5 seconds with > 90% humidity and plunge-frozen into liquid ethane cooled by liquid nitrogen using Vitroblot (FEI Co.)

#### PhuZ-GDP cryo-EM sample preparation

The sample was prepared following the protocol for the PhuZ-GMPCPP sample, but using 80 μM PhuZ in BRB80 pH7.2 and 400 μM GTP. The protein was polymerized for 150 seconds at room temperature. The polymerization time was determined by the right-angle light scattering procedure as described in the Results section.

### Instrumentation and data acquisition

Image stacks were collected on an FEI TF30 Polara microscope (FEI) equipped with a field emission electron source and operated at an accelerating voltage of 300 kV. The images were recorded by the Gatan K2 Summit camera operated in super-resolution counting mode, using UCSFImage4, a semi-automated acquisition package. The images were recorded at 31,000X magnification, corresponding to a super resolution pixel size of 0.61 Å per pixel. The dose rate was set to 10.4 electrons per physical pixel per second on camera and total exposure time was 6 seconds, corresponding to a total accumulated dose of 42 electrons per Å^2^. The images were collected as movie stacks of 30 subframes with 0.2 seconds of exposure time per subframe and acquired over a defocus range from -0.9 μm to -2.3 μm. ~820 image stacks were collected on the PhuZ polymerized in GMPCPP and ~1960 image stacks were collected on the PhuZ polymerized in GTP.

### Image processing

Image stacks were binned twice, which corresponds to a pixel size of 1.22 Å/pixel. Images were motion–corrected using the entire subframes(Li et al., 2013a)Image sums consisting of all 30 subframes were used for processing. Particles were manually selected using boxer in EMAN program suite (Ludtke et al., 1999) 220-pixel filament segments with 40 pixels shift for each segment were extracted from the micrographs. CTFFIND3 was used to determine CTF parameters(Mindell and Grigorieff, 2003). Segments with a low signal-to-noise ratio, bundles or broken segments were discarded after 100 rounds of iterative 2D classification with the Relion software package (regularization parameter T=2) (Scheres, 2012) Segments sorted into highly populated 2D classes were used for further processing. The reconstruction was determined by iterative helical real space reconstruction (IHRSR), essentially as described by Egelman et al., 2000, but following the “gold standard” procedure (Scheres and Chen, 2012) as described in Zehr et al., 2014. SPIDER (Frank et al., 1996) was used for multireference alignment, projection matching, back projection and hsearch_lorentz routine was used for symmetry search (Egelman, 2000). Contrast transfer function (CTF) was corrected by applying Weiner filter to the entire micrograph using Focusramp program (written by D.A.A.). Next, refinement of the alignment parameters together with 3D classification were performed with FREALIGN (Grigorieff, 2007; Lyumkis et al., 2013). Initially, 3D maps were refined against a single reference until no further improvements in resolution estimations were observed. Data up to 10Å were included in the initial rounds of the refinement and up to 5 Å for the PhuZ-GMPCPP map and PhuZ-GDP class 1 in the final rounds. PhuZ-GMPCPP and PhuZ-GDP data sets were divided into 3 classes using RSAMPLE and classified. Refinement iterations were performed until the segment composition for each class and occupancy for each segment converged to nearly constant values. For each of the classes, segments with occupancies of > 70% were extracted and further refined against a single reference. 1 major class was obtained for PhuZ-GMPCPP and three for PhuZ-GDP. The 3D reconstructions generated by FREALIGN (not shown) were prepared for visualization by applying negative B factors using the bfactor program (http://grigoriefflab.janelia.org/bfactor) and were further refined using Frealix (Rohou and Grigorieff, 2014).

For all datasets, Frealix was operated in its “single-particle” mode, wherein continuity restraints along filaments are dropped and segments are treated as mostly independent objects, with the exception that filament polarity is enforced so that segments belonging to the same filament must have similar orientations.

#### GMPCPP dataset refinement

An initial reconstruction by Frealix using alignment parameters from Frealign had an estimated resolution (FSC = 0.143) of 4.9. Initial refinement (using frequencies up to 1/8 Å^-1^) consisted mostly of fixing inconsistencies in polarity among waypoints from the same filament. The high-resolution cutoff was gradually increased to 6 Å (over 25 rounds of refinement), at which point the estimated resolution was 4.0 Å and the FSC at 5.0 Å was 0.79 (compared to 0.2 before Frealix refinement).

Frealix was then modified to accept raw, unaligned frames as inputs (rather than averages of aligned frames). This allowed us to estimate local in-plane shift parameters for each filament waypoint at each frame, using an implementation of the unblur (Grant and Grigorieff, 2015) algorithm on ~ 1,500 × 1,500 Å sub-frames centered around each filament waypoint. After this subframe alignment, waypoint parameters (orientations, shifts) were further refined over 25 iterations during which the length of segments (centered at each waypoint) was decreased in 50-Å steps from 300 Å to 150 Å and then kept constant, and the refinement cutoff raised from 1/6 Å-1 to 1/5 Å-1. This gave a map with an estimated resolution of 3.7 Å and an FSC at 4.5 Å of 0.82. Finally, 3 cycles of refinement were conducted where each cycle included 20 randomized start of the optimizer. The final reconstruction included frames 1 to 30 (total exposure 42 electrons/Å^2^), weighted by an isotropic exposure filter (Grant and Grigorieff, 2015) and an anisotropic drift filter reflecting the directional attenuation of signal due to specimen motion (Frank, 1969) at each waypoint, though the latter did not seem to significantly affect the quality of the reconstruction. The final overall resolution estimate was 3.5 Å, with an FSC of 0.83 at 4.5 Å. At several times during processing, we attempted selecting subsets of frames for reconstruction and refinement (e.g. frames 1-14 or 3-14). The final reconstruction was not improved by such selections, relative to using the full, dose-filtered, stack of frames.

#### Estimation of local helical parameters

A new Frealix feature was developed to enable the refinement of local helical parameters by the same minimizer, and as part of the same scoring function, as other waypoint parameters (shifts and orientation). To this end a new forward-projection method was implemented to compute projections from an input 3D map, whereby the central helical asymmetric unit of the input 3D volume (the boundaries of which are defined as in Figure 4 of Rohou & Grigorieff 2014) is masked and real-space projected *N* times, where *N* is the number of helical asymmetric units present in the 2D projection, with appropriate Euler angles and translational shifts for each of the asymmetric units as predicted by the local helical rise and twist. The resulting composite projections are equivalent to projections obtained following helical symmetrization of the whole 3D volume but are obtained much more quickly. Using this new way of generating composite projection makes it possible to vary helical parameters at each filament waypoint, under control of the optimizer, simultaneously (if desired) with the other refinement parameters. We used this to refine helical parameters for all waypoints in all our datasets. Based on the resulting per-waypoints helical parameters, we were then able to select waypoints within a tighter range of helical parameters. In the case of the GMPCPP dataset, selecting ~61% of helical subunits (see Table 2) led to a slight but noticeable deterioration of the map overall (with possibly some local improvements), so the final map used for interpretation included all subunits. For the GDP1 dataset, selecting 61% subunits in the same manner (Table 2) led to a significantly improved map (although the FSC was not appreciably increased), so we chose to interpret that map.

#### GDP dataset refinement

Other than the selection of subunits based on local helical parameters as summarized in Table 2, refinement with Frealix did not give appreciable improvements in map quality when compared to the maps obtained by FREALIGN. The final refinement of the GDP1 dataset used data up to 1/5 Å^-1^ and a segment length (around each waypoint) of 200 Å.

### Model building and refinement

Because the resolution of the map is not uniform and is somewhat lower at the outer surface of the filament, we combined several strategies to obtain a chemically accurate model. First, a homology model of PhuZ from the closely related bacteriophage ՓPA3 crystal structure was adjusted by flexible fitting with MDFF (Trabuco et al., 2008), greatly facilitating the necessary rebuilding of the backbone and side-chains in some areas and providing general stereochemical correctness. As necessary, minor manual adjustments to the backbone and side-chain atoms were made using Coot(Emsley and Cowtan, 2004), followed by refinement of coordinates and B factors were in REFMAC(Brown et al., 2015). Finally, MDFF was again used to optimize local stereochemistry, especially for distal parts of the side chains and side chains not resolved in the cryo-EM maps The individual model-versus-map FSC curves indicate that the model agrees better with the map using the combined approach than that of MDFF alone (Fig. S1B). Statistics for the model are listed in the Table S2, indicating good stereochemistry, model correctness via their EMRinger scores(Barad et al., 2015). The score improved from 2.2 for the model generated by MDFF alone to 2.7 when the combined approach was used (Table S2). Both scores were substantially above 1.0, the threshold value for models fitted into EM maps resolved in the range from 3 Å to 4 Å, attesting to their high quality.

#### PhuZ-GMPCPP model building

An initial homology model of PhuZ from the bacteriophage ՓPA3 was built using MODELLER software and the X-ray structure of PhuZ from the bacteriophage 201Փ2-1 (PDB ID:3r4v) as a template(Kraemer et al., 2012; Sali and Blundell, 1993). The helical symmetry parameters were applied to the model to built a starting model of PhuZ filament consisting of 9 subunits, and the model was rigid-body fitted into the cryo-EM density using Situs software. The filament model was flexibly fitted into the cryo-EM density using MDFF (Trabuco et al., 2008). A central subunit, refined in the presence of neighboring subunits, was extracted for further model building. To improve the fit of backbone atoms and side, real space refinement of the model was performed with Coot software (Emsley and Cowtan, 2004). After one round of model building, the resulting model was used to build a nonamer again. For the first round of refinement with REFMAC v.5.8 secondary structure restraints were generated using ProSMART. A section of the cryo-EM density, corresponding to the model, was masked out and converted to structure factors using “SFCALC”mode in CCP4. The structure factors were used for restrained refinement of the filament model. The refinement included data up to 3.5 Å. Iterative rebuilding of the model using Coot and REFMAC v.5.8 was repeated until no further improvements in the R-factor and Fourier shell correlation of the model to the map. The stereochemical quality of the final structure was validated by PROCHECK.

### Structure visualization and analysis

Molecular graphics and analyses were performed with UCSF Chimera (Pettersen et al., 2004). Alignment of protein structures was done using FATCAT software(Ye and Godzik, 2003). The difference map was generated using diffmap program (http://grigoriefflab.janelia.org/diffmap).

## CONTRIBUTIONS

E.A.Z. purified protein samples and determined conditions for imaging. E.A.Z. and K.A.V. carried out data acquisition. E.A.Z. and A.R. carried out image processing and analysis. E.A.Z. and Y.L. built atomic models. All authors contributed to experimental design and manuscript preparation.

## ACKNOWLEDEMENTS

We thank Sam Li and Axel Brilot and for invaluable discussions on image processing. Thomas Tomasiak, Alan Brown and Garib Murshudov for help with refining the atomic model with Coot and REFMAC. This work was supported by HHMI (DAA and NG) and NIH grants GM031627, R35GM118099 (DAA), GM073898 (JP), and GM104556 (JP and DAA).

